# Autophagy protects pancreatic β-cells during hypoxia and islet transplantation but is compromised by TFEB-lysosomal dysfunction

**DOI:** 10.64898/2026.07.02.736213

**Authors:** Yuanjie Zou, Daniel J. Pasula, Renmei Tang, Mitsuhiro Komba, Derek L. Dai, Galina Soukhatcheva, C. Bruce Verchere, Dan S. Luciani

## Abstract

Hypoxia is a potent stressor and a major cause of β-cell failure and loss after islet transplantation. Autophagy is a critical homeostatic mechanism that preserves organelle integrity and metabolic balance in cells under stress, but whether it supports β-cell adaptation to sustained oxygen deprivation is unclear. Here, we used β-cell–specific *Atg5* knockout together with hypoxia and transplantation models, to demonstrate that autophagy is a major determinant of β-cell survival during oxygen limitation and supports islet graft function. However, prolonged hypoxia suppressed autophagic flux, reduced lysosomal activity, and led to autophagosome accumulation, indicating failure of the lysosomal clearance pathway. This was accompanied by a marked reduction in transcription factor EB (TFEB) and its lysosomal target genes. Genetic and pharmacological activation of TFEB restored lysosomal gene expression and cathepsin B activity and improved β-cell viability under hypoxia, implicating TFEB decline as a contributor to autophagy–lysosome dysfunction. Together, these findings outline a sequence in which autophagy initially safeguards β-cells but becomes ineffective under sustained hypoxia as TFEB levels fall, identifying TFEB as a potential target to strengthen β-cell resilience and survival in islet transplantation.

## INTRODUCTION

Pancreatic β-cells rely on mitochondrial oxidative metabolism to sustain their energy demands [1]. In islet transplantation, delayed revascularization exposes grafts to oxygen tensions as low as 1%, levels sufficient to impair insulin secretion and trigger apoptosis [2–4]. Although islet replacement is an effective therapy for type 1 diabetes (T1D), early post-transplant β-cell loss continues to hinder long-term success [5]. Because stem cell-derived islets experience similar ischemic stress before vascular integration [6], understanding how β-cells adapt, or fail to adapt, to hypoxia is essential for improving the survival of both donor- and stem cell-derived islet grafts.

To cope with metabolic and oxidative stress, β-cells engage multiple adaptive responses, including antioxidant defenses, the unfolded-protein response (UPR), and autophagy, which together help maintain cellular homeostasis. Autophagy functions as a cellular clearance pathway that removes damaged organelles and misfolded proteins, thereby supporting β-cell survival and insulin secretory function [7]. However, it remains unclear whether autophagy is protective or detrimental for β-cells under chronic hypoxia. In other cell types, hypoxia-induced autophagy can either promote survival or contribute to cell death depending on the intensity and duration of stress [8]. While studies in β-cell lines have shown beneficial effects of autophagy activation under intermittent hypoxia [9], the role of autophagy during sustained oxygen deprivation or in the context of islet transplantation has not been established.

To address this gap, we used mouse models with both in vivo and in vitro β-cell *Atg5* knockout, which disrupts autophagosome formation, to examine how autophagy and related stress pathways affect β-cell survival under sustained hypoxia and in syngeneic islet transplantation. Our results demonstrate that autophagy protects β-cells in both contexts. We further assessed whether this adaptive process remains effective during prolonged hypoxia and investigated the underlying regulatory mechanisms. We found that chronic oxygen deprivation impairs autophagic flux, reduces lysosomal gene expression, and disrupts transcriptional regulation by transcription factor EB (TFEB). Collectively, these findings reveal that hypoxic stress undermines a key β-cell survival pathway and suggest that TFEB loss may contribute to lysosomal and autophagic dysfunction relevant to islet transplantation and diabetes.

## METHODS

### Mice

The following animals were originally purchased from The Jackson Laboratory (Bar Harbor, ME, USA) and then further bred in-house: C57BL/6J (#000664), C57BL/6-Tg(CAG-RFP/EGFP/Map1lc3b)1Hill/J (common name: CAG-RFP-EGFP-LC3, #027139) and Gt(ROSA)26Sor*^tm4(ACTB-tdTomato,-EGFP)Luo/^*J (common name: mTmG, #007576). B6.129S-Atg5*^tm1Myok^* mice (common name: *Atg5^flox/flox^*) were kindly gifted from Dr. Noboru Mizushima (via RIKEN) [10]. All mice were housed in groups of 2-4 per cage in a temperature-controlled conventional animal facility (22 ± 2°C) with 40-60% relative humidity and a 12-hour light/12-hour dark cycle. Animals had ad libitum access to water and standard rodent chow (18%Kcal^Fat^, 24%Kcal^Protein^, 58%Kcal^Carbohydrate^; #2918, Envigo, Indianapolis, IN, USA). Bedding and environmental enrichment were provided, and cages were changed according to institutional standard operating procedures. All animal procedures approved by the University of British Columbia Animal Care Committee and conducted in accordance with institutional, national and international guidelines.

### Islet isolation, culture and dispersion

After anesthesia and euthanasia, mouse pancreatic islets were isolated by a collagenase digestion/filtration method as described before [11, 12]. In brief, approximately 2 ml of collagenase (1000 U/mL, #C7657; Sigma-Aldrich) in Hank’s solution (138 mM NaCl, 5.3 mM KCl, 0.34 mM NaH_2_PO_4_, 0.44 mM KH_2_PO_4_ and 5.56 mM glucose) was administered via the pancreatic duct. The pancreas was removed and incubated in collagenase solution at 37°C for 15 min and then shaken by hand to break down the connective tissue and release the islets. The dispersed tissue was then washed with Hank’s solution containing 1 mM CaCl_2_. After centrifugation for 1 min at 200g and filtration through a 70 µm nylon cell strainer (#352350 Falcon), the islets were re-suspended in Roswell Park Memorial Institute 1640 media (RPMI, #11875-093, Thermo Fisher Scientific, Waltham, MA, USA) and picked by hand for purification. Unless otherwise indicated, mouse islets were cultured at 37°C in 5% CO_2_ in RPMI completed with 10% fetal bovine serum (FBS, #12483-020, Thermo Fisher Scientific) and 2% Penicillin-Streptomycin (#15140-122, Thermo Fisher Scientific). For single cell studies, mouse islets were dispersed with 0.01% trypsin (#25300-054, Thermo Fisher Scientific) onto glass coverslips (#16004-310, VWR international, LLC, Radnor, PA, USA) or ibidi 8-well chamber-slides with polymer (#80806/#80826, ibidi, Munich, Germany) or glass bottom (#80827, ibidi). Islet cells were cultured at 37°C in 5% CO_2_ in complete RPMI 1640 media for 1-2 days after dispersion before staining or imaging. Human islets were provided by Clinical Islet Laboratory through the Alberta Islet Distribution Program. The islets were purified by hand-picking and then dispersed onto ibidi 8-well chamber slides (#80806/#80826, ibidi) for microscopy. The dispersed islet cells were cultured in CMRL media (#11530037, Thermo Fisher Scientific) completed with 10% FBS, 2% Penicillin-Streptomycin, at 22°C in 5% CO_2_ for 1-2 days before experiments.

### Adenoviral transduction

Dispersed islet cells were allowed to recover for 1-2 days in ibidi chamber slides before being incubated in completed RPMI 1640 media containing adenovirus at the indicated MOI (viral particles added per cell). Following overnight transduction, the media was replaced by fresh virus-free RPMI, and the cells were cultured for additional 3-4 days before being used for imaging or RNA analysis.

### Glucose metabolism

Non-fasting body weight and blood glucose measurements were recorded in the morning (9 am-10 am), using OneTouch Ultra® or OneTouch Verio Flex® blood glucose meters and test strips (LifeScan Inc., Malvern, PA, USA). Glucose tolerance was assessed by intraperitoneal glucose tolerance tests (IPGTT). IPGTTs were performed in the afternoon, starting at 3-4 pm, after a 6 hour fast. Unless otherwise indicated, mice were injected with 2 g glucose/kg body weight. Blood glucose was measured at 0, 15, 30, 60, 90 and 120 min after glucose injection.

### Islet Transplantation

A marginal mass syngeneic islets transplantation model was used to study the role of autophagy under post-transplantation stresses that occur independent of autoimmune graft rejection [13]. Female *Atg5^flox/flox^* mice of 10-14 weeks of age were used as transplant donors. Two weeks before islet isolation, double-stranded adeno-associated viruses (dsAAV) encoding Rat Insulin Promoter-1 (RIP1)-driven Cre recombinase (dsAAV6-RIP1-Cre) or empty dsAAV control viruses (dsAAV6-Empty) were delivered into the pancreas through an intraductal injection to create *Atg5* knockout and control donor islets, respectively [14, 15]. After islet isolation, *Atg5* deletion was verified using aliquots of the islets by qPCR. Male *Atg5^flox/flox^* mice of 8-12 weeks of age were injected i.p. with a single high STZ dose (175 mg/kg; #S0130, Sigma-Aldrich, St Louis, MO, USA) to induce diabetes. Four days after injection, mice showing hyperglycemia (two consecutive days with blood glucose >20 mM) were used as islet transplant recipients. A marginal mass of islets was transplanted under the kidney capsule, essentially as described [16]. Donor islets were isolated one day prior, pooled and then hand-picked for equal size grafts on the morning of islet transplantation. Each recipient received a graft containing 300-350 hand-picked islets. The marginal number of islets was established in prior tests on mice of the same sex and age. Graft recipients were monitored for body weight and blood glucose every day for 2 weeks post-surgery, and intraperitoneal glucose tolerance tests (IPGTTs) were performed 2 weeks after the transplant, or at any earlier time if the recipients became hyperglycemic due to graft failure.

### Hypoxic culture

Hypoxic culture (1% O_2_, 5% CO_2_ and 94% N_2_) of intact mouse islets, as well as dispersed mouse and human islet cells for immunofluorescent imaging and lysosomal function assays, was done in an incubator sub-chamber (C-Chamber) with CO_2_ and O_2_ levels maintained by ProCO_2_ 120 and ProOx 110 controllers (Biospherix, Parish NY, USA). After hypoxic culture, whole islets were immediately processed for qPCR or Western blot analysis. For immunostaining, hypoxic islet cells were immediately fixed in 4% paraformaldehyde (PFA, #AC41678, Fisher Scientific). For imaging-based analysis of cell survival and autophagic flux (detailed below), dispersed islet cells were cultured at 37°C under hypoxia (1% O_2_, 5% CO_2_ and 94% N_2_) or normoxia (20% O_2_, 5% CO_2_ and 75% N_2_) in a Tokai Hit INUBTFP-WSKM microscope stage-top incubator.

### Glucose-stimulated insulin secretion

Fifteen size-matched islets were placed into low-retention microcentrifuge tubes and cultured in Krebs-Ringer buffer (KRB) with 3 mM glucose for 1 hour prior to the experiments. The islets were then transferred to 300 μl fresh KRB containing 3 mM glucose, 15 mM glucose, each for 1 hour sequentially. The supernatants from each stage were collected and assayed for secreted insulin using a Mouse Ultrasensitive Insulin ELISA (#80-INSMSU-E10, ALPCO, Salem, NH, USA) following the manufacturer’s instructions.

### Cell Death

For quantification of cell death, mouse islet cells were dispersed and seeded onto 25 mm glass coverslips or ibidi 8-well chamber-slides (#80806/#80826). The islet cells were pre-loaded for 30 min with 0.05 μg/ml Hoechst 33342 (#H3570, Thermo Fisher Scientific) and 0.5 μg/ml propidium iodide (PI, #P1304MP, Thermo Fisher Scientific). To quantify cell death, the cells were imaged during culture in normoxia or hypoxia within a Tokai Hit INUBTFP-WSKM Stage-top incubator at 37℃ and images were taken using the wide-field setting on a Leica DMI6000 inverted microscope using a Leica HCX Plan FLUOTAR L 20x objective (#506242), DAPI ET and DsRed ET filter cubes (#49005, Chroma). The percentage of dead cells (PI^+^ / Hoechst^+^) was quantified using ImageJ.

### Cytosolic Ca^2+^ imaging

Cytosolic Ca^2+^ was imaged in intact islets using Fura-2 AM (#F1221, Thermo Fisher Scientific) [11]. After isolation, 10-15 islets were picked and seeded onto a 25 mm glass coverslip (#16004-310, VWR) placed in 35 mm culture dishes (#353001, Corning Incorporated, Corning, NY, USA). Islets were incubated for 3 days to attach before imaging. Islets were loaded with 5 µM Fura-2 for 30 min and then perifused with a modified Ringer’s Buffer (5.5 mM KCl, 2 mM CaCl_2_, 1 mM MgCl_2_, 20 mM HEPES, 144 mM NaCl) containing 3 mM glucose for at least 30 min to reach a stable baseline. Glucose responsiveness was tested by exposing the islets to a ramp of Ringer’s containing 3 mM, 5 mM, 10 mM and 15 mM glucose. Fura-2 was excited at 340 nm and 380 nm and the emission was collected at 502-538 nm every 10 seconds in wide-field settings on a Leica DMI6000 inverted microscope (Leica, Wetzlar, Germany) equipped with a Leica HCX Plan FLUOTAR L 10x objective (#506505), Fura-2 filter cube (#79001, Chroma, Bellows Falls, VT, USA).

### Cathepsin B activity

Lysosomal cathepsin B activity was examined in dispersed mouse islet cells using the Magic Red^®^ assay Kit (#ICT937, Bio-Rad), following the manufacturer’s guidelines. To minimize effects of reoxygenation, islet cells were briefly fixed with 0.5% PFA for 30 min before staining. Widefield images were taken using a Leica HCX Plan FLUOTAR L 10x objective (#506505) and DsRed ET filter cube (#49005, Chroma). Magic Red fluorescence intensities were quantified in ImageJ.

### 3D confocal imaging and quantification of autophagic flux

Dispersed islet cells from CAG-RFP-EGFP-LC3 mice were seeded in ibidi 8-well glass bottom chamber slides coated with 804G cell-derived extracellular matrix to improve islet cell attachment and spreading [17]. The islet cells were imaged after 24 hours culture in a Tokai Hit INUBTFP-WSKM stage-top incubator under normoxia or hypoxia. Stack images were taken using the high-speed (8,000 Hz) resonance scanner of a Leica SP8 Laser Scanning Confocal Microscope and collected using an HC PL APO oil 63x objective. Signals were collected at the spectrum of EGFP (excitation: 488 nm, emission: 493-550 nm) and mRFP (excitation: 561 nm, emission 650-765 nm). The resulting image stacks were deconvolved in Huygens Pro® deconvolution wizard and then reconstructed 3-dimensionally in ImageJ using the “3D object count” plugin. Colocalization of GFP- and RFP-positive signal and puncta volumes were quantified using the ImageJ “Coloc 2” and “3D object count” plugins.

### Immunofluorescence staining

Islet cells from C57BL/6J mice or non-diabetic human donors were dispersed onto ibidi 8-well chamber slides and cultured under hypoxia or normoxia with or without additional treatments, as indicated. Cells were fixed in 4% PFA, rinsed with PBS (#L14190-136, Thermo Fisher Scientific), permeabilized using 0.05% Triton X-100 (X100, Sigma-Aldrich) for 5 min, and blocked for 1 hour with Serum Free Protein Block (#X0909, Agilent-DAKO) at room temperature. Primary antibodies were added at 4℃ overnight and secondary antibodies for ∼1 hour at room temperature. Primary and secondary antibodies used for immunostaining are listed in **Table 1**.

**Table 1.**
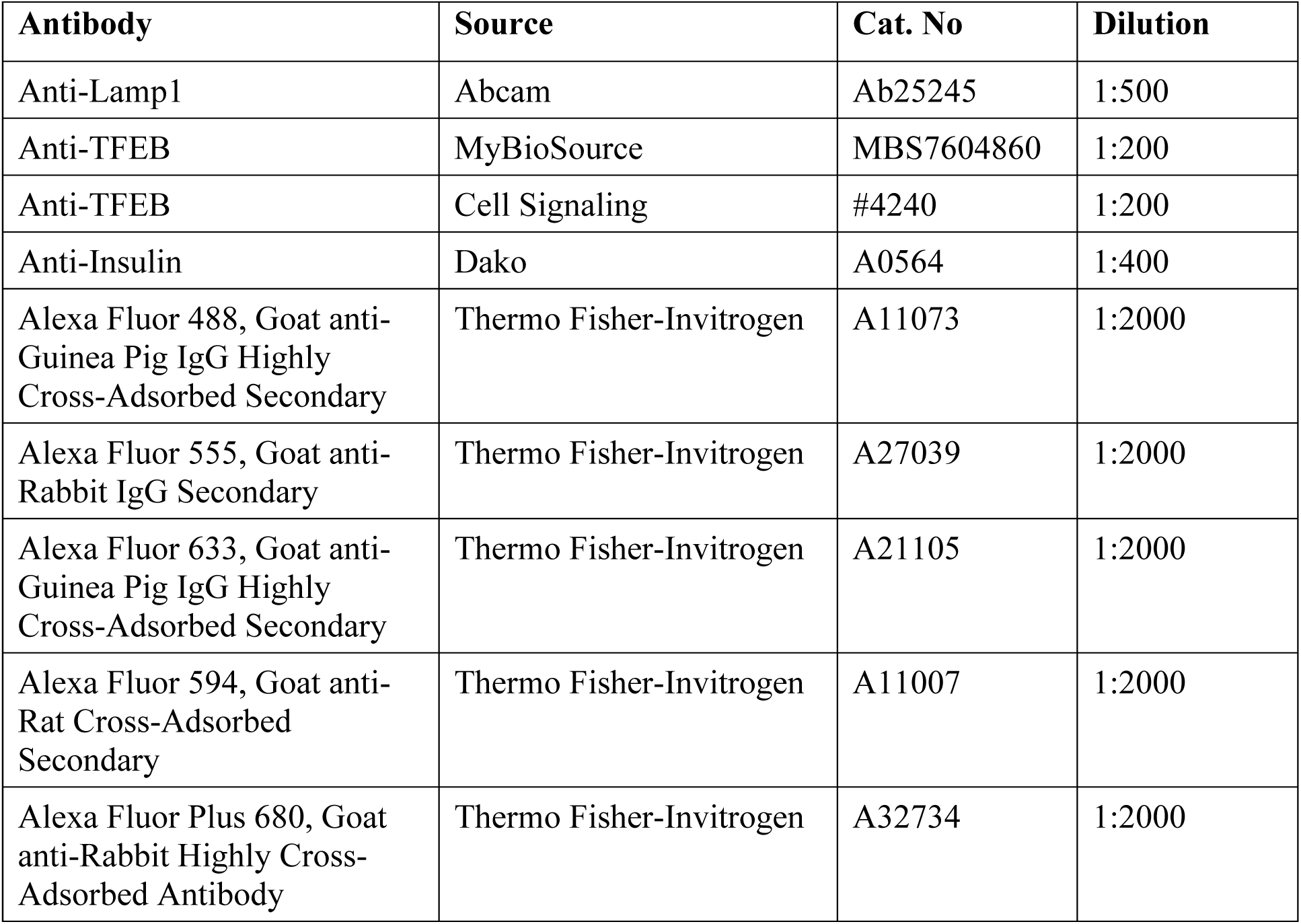
Antibodies used for immunofluorescent staining.

| Antibody | Source | Cat. No | Dilution |
| --- | --- | --- | --- |
| Anti-Lamp1 | Abcam | Ab25245 | 1:500 |
| Anti-TFEB | MyBioSource | MBS7604860 | 1:200 |
| Anti-TFEB | Cell Signaling | #4240 | 1:200 |
| Anti-Insulin | Dako | A0564 | 1:400 |
| Alexa Fluor 488, Goat anti-Guinea Pig IgG Highly Cross-Adsorbed Secondary | Thermo Fisher-Invitrogen | A11073 | 1:2000 |
| Alexa Fluor 555, Goat anti-Rabbit IgG Secondary | Thermo Fisher-Invitrogen | A27039 | 1:2000 |
| Alexa Fluor 633, Goat anti-Guinea Pig IgG Highly Cross-Adsorbed Secondary | Thermo Fisher-Invitrogen | A21105 | 1:2000 |
| Alexa Fluor 594, Goat anti-Rat Cross-Adsorbed Secondary | Thermo Fisher-Invitrogen | A11007 | 1:2000 |
| Alexa Fluor Plus 680, Goat anti-Rabbit Highly Cross-Adsorbed Antibody | Thermo Fisher-Invitrogen | A32734 | 1:2000 |

Immunocytochemistry images were analyzed for their mean intensity and/or puncta counts using ImageJ software. Masks for the cellular nuclear area, cytosolic area and/or the whole cell area were generated using built-in threshold functions. Puncta were identified and analyzed for size and fluorescence intensity using ImageJ Auto Local Threshold and Particle Analysis functions. All imaging settings were held constant across conditions and samples within experiments.

### RT-qPCR

Total RNA was extracted from mouse islets using RNeasy Micro Kit (#74104, Qiagen, Hilden, Germany) according to the manufacturer’s guidelines. RNA yield was measured on a NanoDropTM 2000c (#ND2000CLAPTOP, Thermo Fisher Scientific) spectrophotometer. cDNAs were generated through reverse transcription by qScript cDNA synthesis kits (#95047, Quanta Bioscience, Gaithersburg, MD, USA). Quantitative PCR (qPCR) was performed using PerfeCTa SYBR Green SuperMix (#95053, Quanta Bioscience) in the ViiA7 Real-Time PCR System (Thermo Fisher/Applied Biosystems) with primers listed in **Table 2**.

**Table 2.**
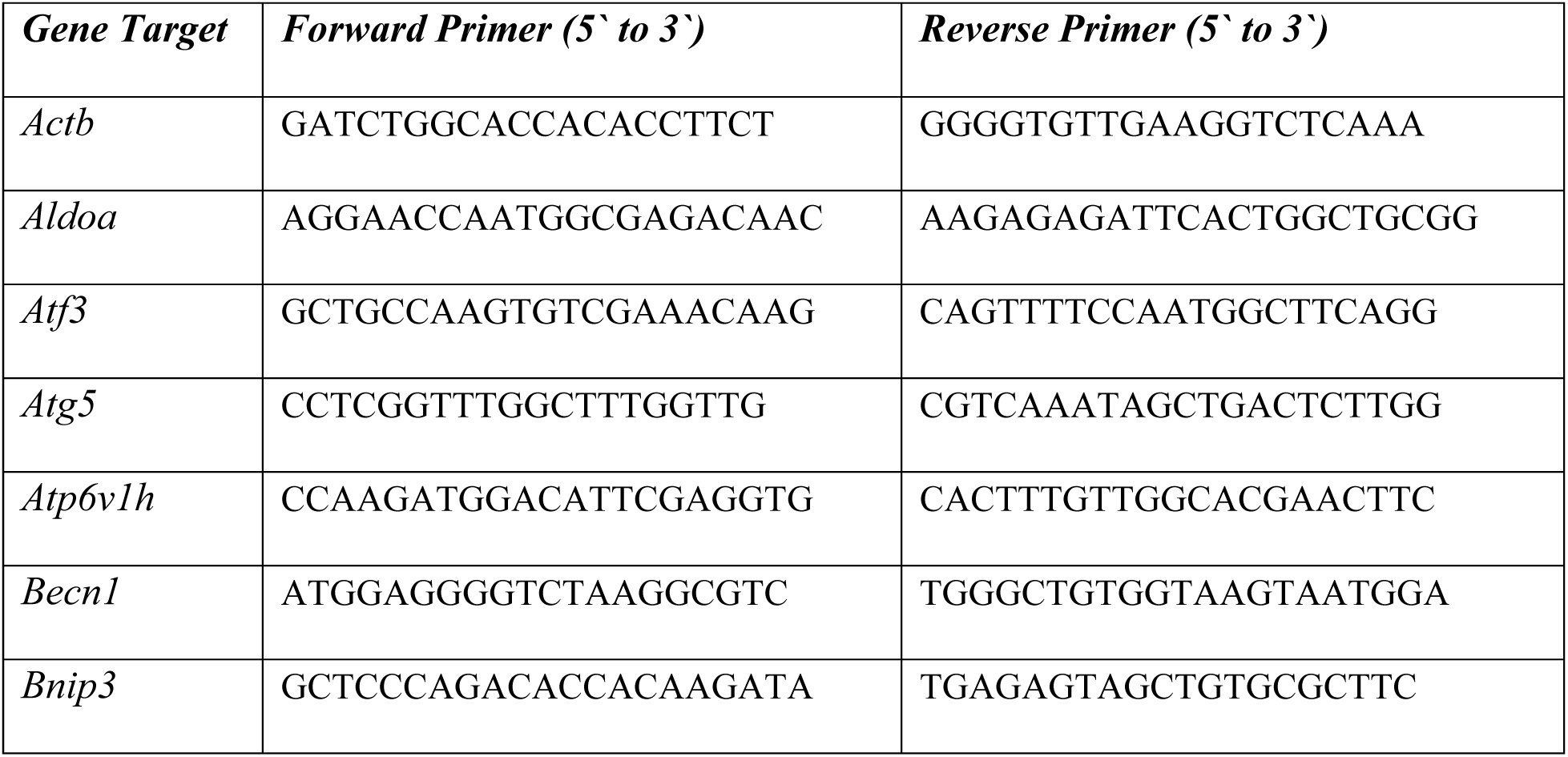
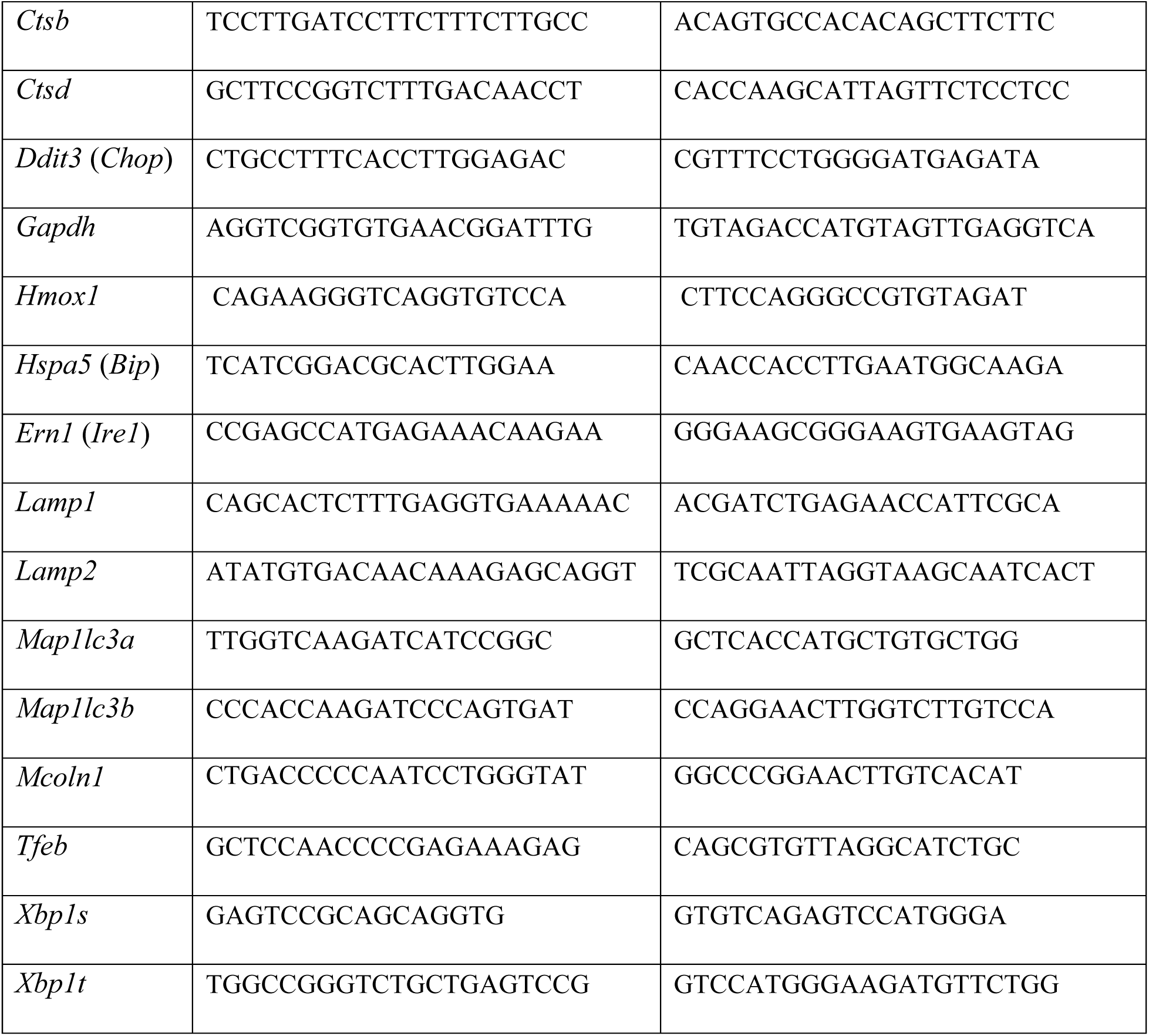
Primers used for SYBR Green qPCR.

### Western blotting

Isolated islets were transferred to a 1.5-ml microcentrifuge tube. After brief centrifugation, culture medium was discarded and 50 μl of RIPA lysis buffer (150 mM NaCl, 50 mM Tris-HCl (pH 8.0), 0.1% Triton X-100, 0.5% sodium deoxycholate, 0.1% SDS, 1 x Halt Protease and Phosphatase inhibitor (#78440, Thermo Fisher Scientific) was added. The cells were lysed by sonication (Misonix S-4000) with 40% power output, 6-10 cycles of 10-sec-ON – 20-sec-OFF. Protein lysates were then separated by 4-20% SDS-PAGE (#4561095, Bio-Rad, Hercules, CA, USA) and transferred to a PVDF membrane (#1620177, Bio-Rad). The membrane was blocked with 5% non-fat dry milk in TBST and incubated overnight with primary antibody at 4°C. After three washes with TBST, the membrane was incubated with HRP-conjugated secondary antibody at room temperature for 1 hour. After three further washes with TBST, the membrane was then incubated with SuperSignal West Pico or Femto ECL Substrates (#34579/#34094, Thermo Fisher Scientific) and the chemiluminescent signals were detected by X-ray film. Antibodies used for western blot are listed in **Table 3**.

**Table 3.** Antibodies used for western blot.

| Antibody | Source | Cat. No | Dilution |
| --- | --- | --- | --- |
| Anti-Beta Actin | Novus Biologicals | NB600-501 | 1:10000 |
| Anti-Atg5 | Cell Signaling | #12994 | 1:1000 |
| Anti-Cre | Novagen | 60950-3 | 1:10000 |
| Anti-LC3 | Novus Biologicals | NB100-2220 | 1:900 |
| Anti-TFEB | Bethyl Laboratories | A303-673A | 1:10000 |
| Anti-mouse IgG<br>HRP-linked Antibody | Cell Signaling | #7076 | 1:3000 |
| Anti-Rabbit IgG<br>HRP-linked Antibody | Cell Signaling | #7074 | 1:3000 |

### Statistical analysis

Data are represented as mean ± SEM and were analyzed in GraphPad Prism 9.0 (La Jolla, California) using Student’s t-test, one-way ANOVA, or two-way ANOVA followed by multiple comparison tests, as appropriate. Statistical significance was set at p < 0.05.

## RESULTS

### Hypoxia leads to loss of adaptive UPR- and autophagy-related transcripts in pancreatic β-cells

To investigate how hypoxia affects pancreatic islets, we cultured C57BL/6J mouse islets under hypoxia (1% O_2_) or normoxia (20% O_2_) and quantified the time-dependent effects on gene expression by qPCR. As expected, known hypoxia-induced transcripts, including *Hmox1*, *Aldoa*, *Gapdh* and *Bnip3*, were rapidly upregulated in hypoxic culture (**Figure 1a**). Hypoxia also significantly induced the expression of pro-apoptotic ER stress-induced genes *Ddit3* (i.e. *Chop*) and *Atf3*, whereas the expression of *Xbp1s* and *Hspa5* (i.e. *Bip*), which are part of the adaptive unfolded protein response (UPR), were reduced or unchanged (**Figure 1b**). These findings agree with a previous report of selective hypoxia-induced impairment of the adaptive UPR [18].

**Figure 1.**
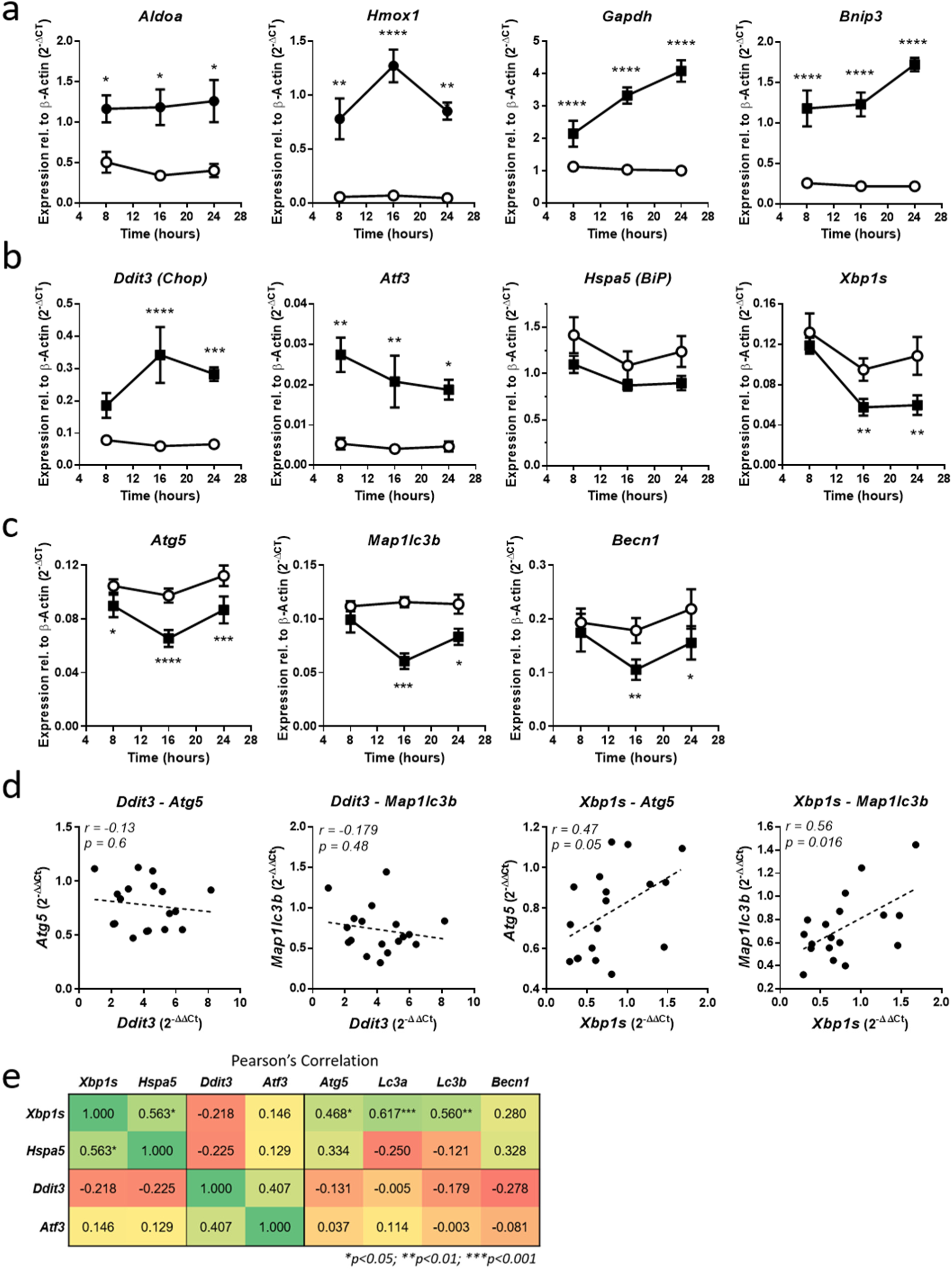
Time-dependent changes to islet ER stress- and autophagy-related gene expression in response to hypoxia. a) Time-dependent expression of known hypoxia-responsive transcripts in islets under normal and 1% O_2_ culture. b) Hypoxia-induced changes to the expression of select islet UPR / ER stress response genes. c) Hypoxia-induced changes to the expression of select autophagy-related genes. d) Examples of linear regression fits and correlation analysis of hypoxia-induced changes to gene expression across all time-points examined in panels (a) & (b). e) Summary of correlation between the expression of UPR transcripts (adaptive *Xbp1s*, *Hspa5*; maladaptive *Ddit3*, *Atf3*) and autophagy-related transcripts (*Atg5*, *Map1lc3a*, *Map1lc3b*, *Becn1*, *Bnip3*). Numbers in the table indicate the Pearson’s correlation coefficient and the statistical significance is indicated by asterisks Data in panels (a)-(c) show the mean ± SEM of 4-6 independent experiments and were analyzed using Two-way ANOVA followed by Tukey’s multiple comparison test. Panels (d) & (e) were analyzed using Pearson Correlation. \**p* < 0.05, \*\**p* < 0.01, \*\*\**p* < 0.001.

Autophagy is well-known for its protective role in β-cells under various stress conditions, including glucolipotoxicity [19, 20] and ER stress [21], but the impact of hypoxia on β-cell autophagy pathways is not clear. To explore this, we quantified the mRNA levels of the key autophagy-regulators *Atg5*, *Map1lc3b* and *Becn1* and found that these were downregulated similarly to the adaptive UPR transcripts, suggesting that autophagy may also be negatively affected by hypoxic stress (**Figure 1c**). To investigate the relationship between the UPR and autophagy, we performed correlation analysis of the changes to gene expression across all time points. This showed that the hypoxia-induced changes to autophagy-related transcripts correlated with changes to the adaptive UPR transcripts, but not to the pro-apoptotic ER stress genes *Ddit3* and *Atf3* (**Figures 1d,e**). Together these results highlight that gene expression in the adaptive UPR and autophagy pathways are correlated and similarly suppressed by hypoxia in mouse islets.

### Autophagy is important for β-cell survival under hypoxia-induced stress

*Atg5* is one of the autophagy-related mRNAs that we found to be down-regulated by hypoxia. It is a key player in the formation of autophagosomes and lack of Atg5 leads to cellular autophagy-deficiency [22]. To investigate the role of autophagy in β-cells under hypoxia, we used adenovirus-mediated expression of Cre recombinase (Ad.Cre) to delete *Atg5* in dispersed islet cells from *Atg5*^flox/flox^ mice. At an MOI of 40, Cre transduction caused recombination in nearly all dispersed islet cells from ROSA^mTmG^ reporter mice (**Figure 2a and Supplemental Fig. 1**) and resulted in ∼90% loss of *Atg*5 mRNA expression in *Atg5*^flox/flox^ islet cells (**Figure 2b**). Comparing hypoxia-induced cell death 4 days after transduction with either control adenovirus (*Atg5*^Ad.Ctrl^) or adenovirus expressing Cre (*Atg5*^Ad.KO^) revealed that autophagy-deficient *Atg5*^Ad.KO^ cells underwent significantly quicker and more severe death than *Atg5*^Ad.Ctrl^ cells (**Figure 2c-e**). This demonstrates that autophagy is an important pro-survival mechanism for pancreatic β-cells during hypoxic stress.

**Figure 2.**
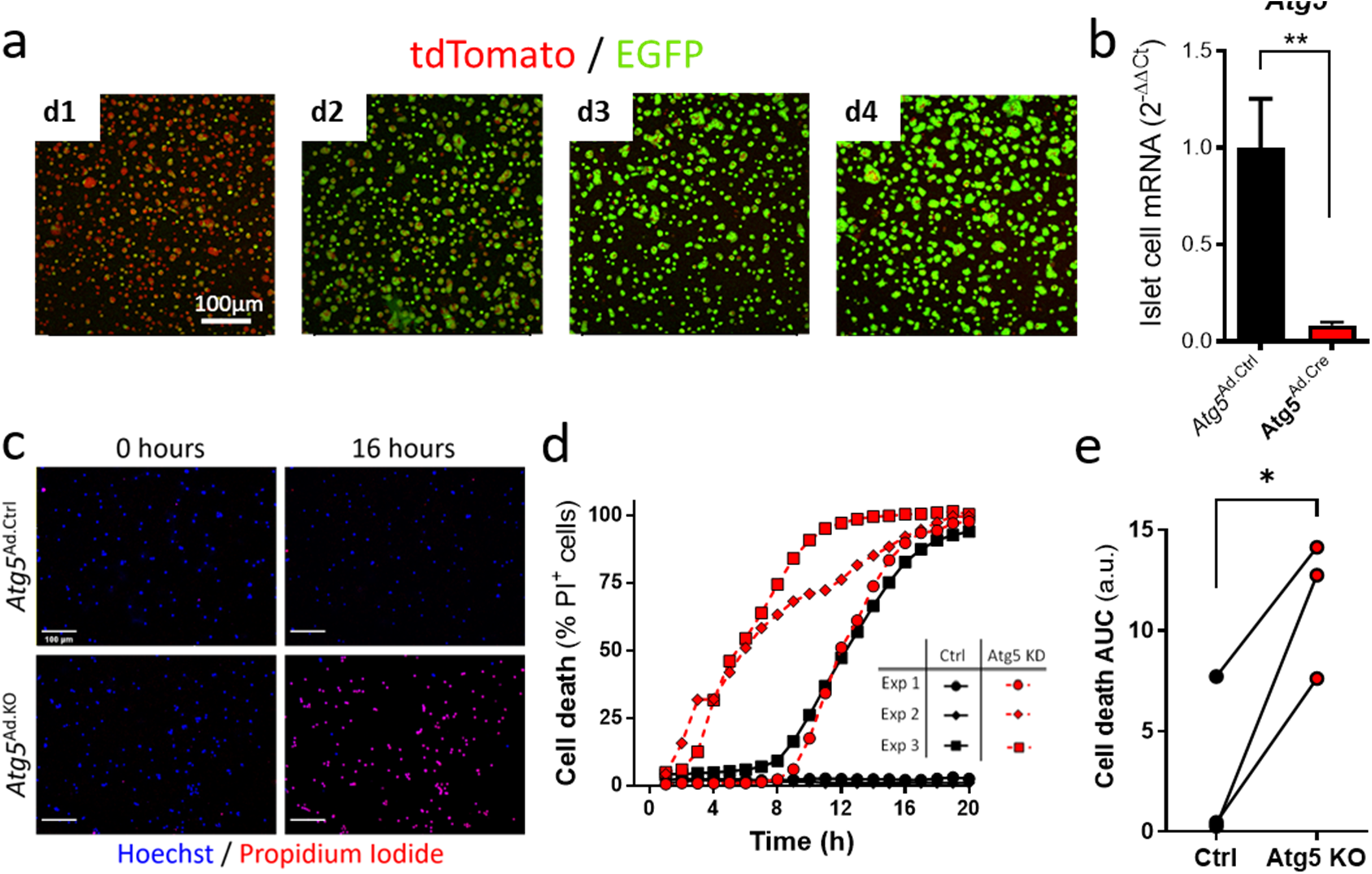
Autophagy is essential for β-cell survival under hypoxia. a) Representative images of dispersed islet cells from ROSA^mTmG^ reporter mice following transduction by adenovirus Cre (Ad.Cre) at MOI 40. Loss of tdTomato and gain of EGFP fluorescence shows the degree of LoxP recombination over 4 days. Reporter activation at different MOIs of Ad.Cre virus are shown in Supplemental Figure 1. b) *Atg5* mRNA levels in *Atg5*^flox/flox^ mouse islet cells after 4 days transduction with MOI 40 Ad.Cre (*Atg5*^Ad.KO^ cells) or Ad.Ctrl (*Atg5*^Ad.Ctrl^ cells) n = 6; \*\**p* < 0.005 by paired t-test. c) Representative images comparing hypoxia-induced cell death (propidium iodide incorporation) of *Atg5*^Ad.KO^ and *Atg5*^Ad.Ctrl^ islet cells at 0 and 16 hrs (scale bar 100 µm). d) Time-course quantification of hypoxia-induced cell death in 3 independent cultures of *Atg5*^Ad.KO^ and *Atg5*^Ad.Ctrl^ islet cells. e) Total cell death (area under the curve; AUC) of each experiment in panel (d). \**p* < 0.05 by Student’s t-test.

### β-Cell autophagy-deficiency accelerates failure of syngeneic islet grafts

We next examined the role of endogenous β-cell autophagy in islet graft function. To do this without confounding physiological defects associated with long-term β-cell autophagy deficiency [23–25], we induced time-controlled *Atg5* knockout by injecting *Atg5*^flox/flox^ mice with dsAAV6 expressing Cre recombinase under the rat *Ins1* promoter (dsAAV6-RIP1-Cre; *Atg5*^AAV.βKO^ mice), or empty dsAAV6 control virus (dsAAV6-Empty; *Atg5*^AAV.Ctrl^ mice). AAVs were injected via the pancreatic duct 2 weeks before the islets were isolated and transplanted into syngeneic recipients with STZ-induced diabetes (**Figure 3a & Supplemental Fig. 2a**). As expected from previous work with this dsAAV6 vector [14, 15], Cre was expressed in insulin-positive β-cells, but not glucagon-positive α-cells (**Figure 3b**). qPCR analysis and western blot confirmed the loss of *Atg5* mRNA and protein in *Atg5*^AAV.βKO^ mouse islets (**Figure 3c**). Reduced protein levels of LC3-II relative to LC3-I indicate that *Atg5*^AAV.βKO^ donor islets had a basal autophagy deficiency, as expected (**Figure 3d**). AAV-induced short-term *Atg5* knockout did not impair β-cell function; *Atg5*^AAV.Ctrl^ and *Atg5*^AAV.βKO^ islet donor mice showed normal glucose tolerance 1 day before islet isolation (**Figure 3e**) and their isolated islets had similar glucose-stimulated Ca^2+^ and insulin secretion responses prior to transplantation (**Figure 3f-h**). After islet transplantation, blood glucose and body weight were monitored in the recipients until the grafts failed or the mice reached our pre-defined endpoint at 14 days post-transplantation (**Figure 3a**). Before and after islet transplantation, body weights did not differ between the two recipient groups (**Supplemental Fig. 2b**) and the transplantation of a marginal mass of islets reversed diabetes in both groups (**Figure 3i**). Importantly, however, autophagy-deficient grafts failed at a faster rate than similar-sized *Atg5*^AAV.Ctrl^ grafts (**Figures 3i,j**). Mice that had received transplants of *Atg5*^AAV.βKO^ islets also had worse glucose tolerance than mice receiving control grafts (**Figure 3k**). At the experimental endpoint, 5 *Atg5*^AAV.Ctrl^ graft recipients and 1 *Atg5*^AAV.βKO^ graft recipient had random blood glucose measurements below the 20 mM cutoff used to define the onset of diabetes. After removal of the graft bearing kidney all mice became diabetic within 4 days, confirming that their normoglycemia was maintained by the transplanted islets (**Supplemental Fig. 2c**). Together, these data show that β-cell autophagy plays an important role in sustaining early islet graft function under the compound hypoxic, inflammatory and glucotoxic stress experienced in syngeneic grafts.

**Figure 3.**
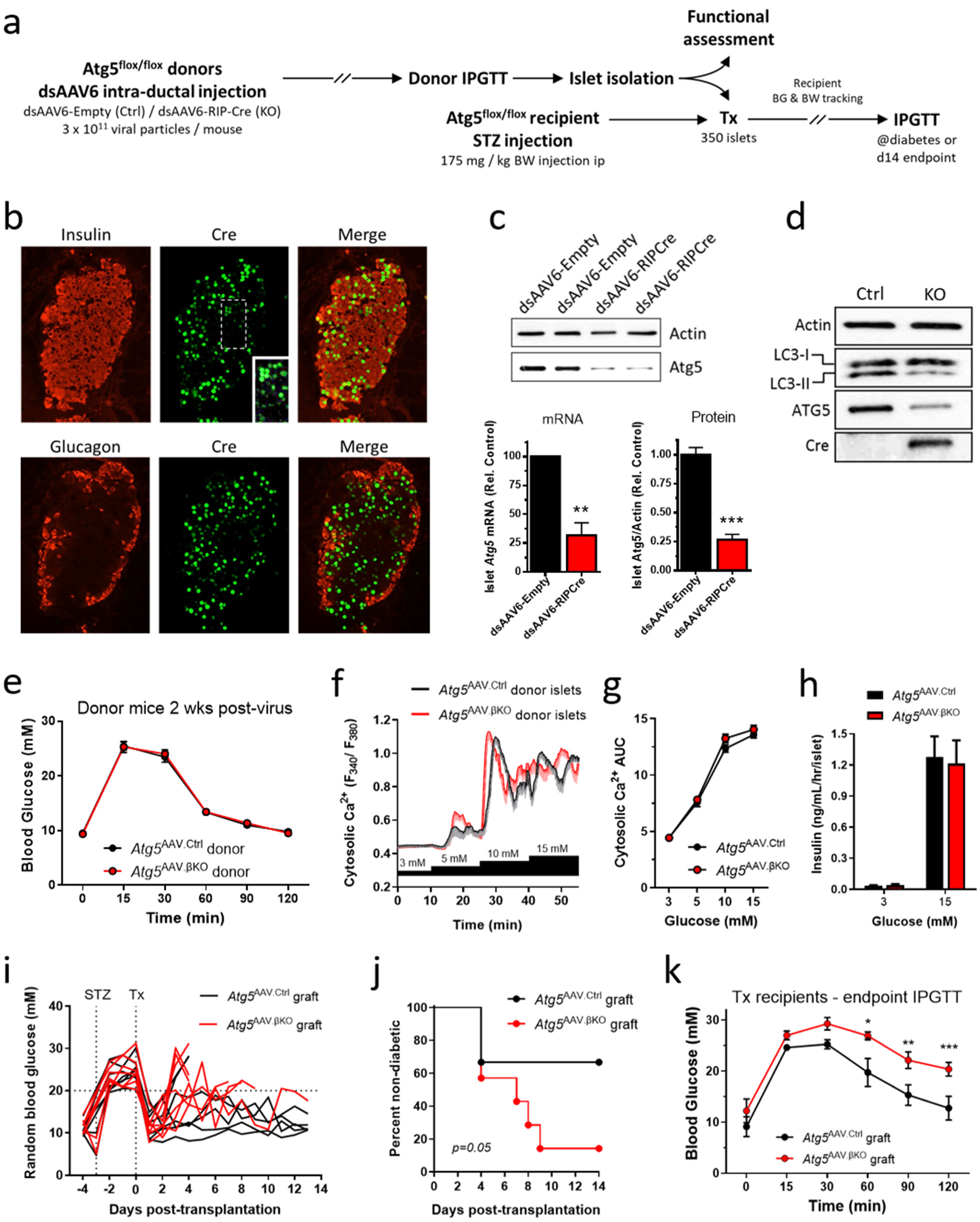
β-Cell autophagy deficiency sensitizes to islet graft failure. a) Schematic outline of workflow for marginal mass islet transplantation experiment. b) Illustrative immunostaining showing Cre expression in β-cells, but not α-cells, 2 weeks after intraductal dsAAV6-RIP1-Cre injection. Insert: Contrast enhancement reveals low levels of Cre expression in seemingly negative β-cells. c) Western blot and qPCR quantification demonstrating loss of *Atg5* mRNA and protein in islets from dsAAV6-RIP1-Cre-injected (*Atg5*^AAV.βKO^) mice compared to islets from mice injected with dsAAV6-Empty virus (*Atg5*^AAV.Ctrl^) (n = 3 islet preparations of each genotype, \*\**p* < 0.01, \*\*\**p* < 0.001 by t-test). d) Western blot showing that islets from Cre-expressing *Atg5*^AAV.βKO^ islets have reduced levels of LC3-II, indicating loss of basal autophagy. e) IPGTT demonstrating that the *Atg^5^*^AAV.βKO^ donor mice had normal glucose tolerance 2 weeks after AAV injection (n=12 mice in each group). f) Fura-2 fluorescence imaging demonstrating similar glucose-stimulated Ca^2+^ responses in aliquots of *Atg5*^AAV.Ctrl^ and *Atg5*^AAV.βKO^ donor islets (14 Ctrl and 13 KO islets from 3 separate and matched donor preparations). g) Area under the curve (AUC) summary of Ca^2+^ imaging results from panel (f). h) Static incubation demonstrating robust and comparable glucose-stimulated insulin secretion in aliquots of *Atg5*^AAV.Ctrl^ and *Atg5*^AAV.βKO^ donor islets (n=4 islet preparations). i) Random blood glucose measurements from individual *Atg5*^flox/flox^ mice made diabetic by STZ and receiving marginal mass islet grafts from *Atg5*^AAV.Ctrl^ or *Atg5*^AAV.βKO^ donors. Endpoint was defined by 2 consecutive days with blood glucose > 20 mM or 2-weeks post-transplantation for mice with no graft failure. j) Survival curve showing diabetes incidence in islet graft recipients (n=7 recipients of each graft genotype). k) Endpoint IPGTT of a subset of graft recipients (n=4 recipients of each graft genotype, \*\**p* < 0.01, \*\*\**p* < 0.001 by repeated measures two-way ANOVA followed by Sidak’s multiple comparison tests).

### Endogenous autophagic flux is impaired in hypoxic β-cells

Given the importance of autophagy for β-cell viability and the impaired expression of autophagy genes under hypoxia, we next examined how endogenous autophagy is affected by hypoxia. For this, we cultured islet cells from RFP-EGFP-LC3 reporter mice [26] (**Figure 4a**) under normoxia and hypoxia and quantified autophagic flux using 3D live-cell imaging. As illustrated in 3D reconstructions of reporter islet cells (**Figure 4b, Supplemental Videos 1-4**) and quantified by EGFP/RFP colocalization analysis (**Figure 4c**), prolonged (24 hours) hypoxia led to an accumulation of autophagosomes. This resembled what was seen when lysosomal function and autophagic flux were inhibited with chloroquine. Comparing the two conditions (**Figure 4c**; 2nd and 3rd bars) shows that late stage autophagic flux is almost completely halted in a large fraction of the hypoxic cells. An increase in the volume of individual autophagosomes and autolysosomes further supports that hypoxia disrupts normal homeostatic control of these organelles (**Figure 4d**).

**Figure 4.**
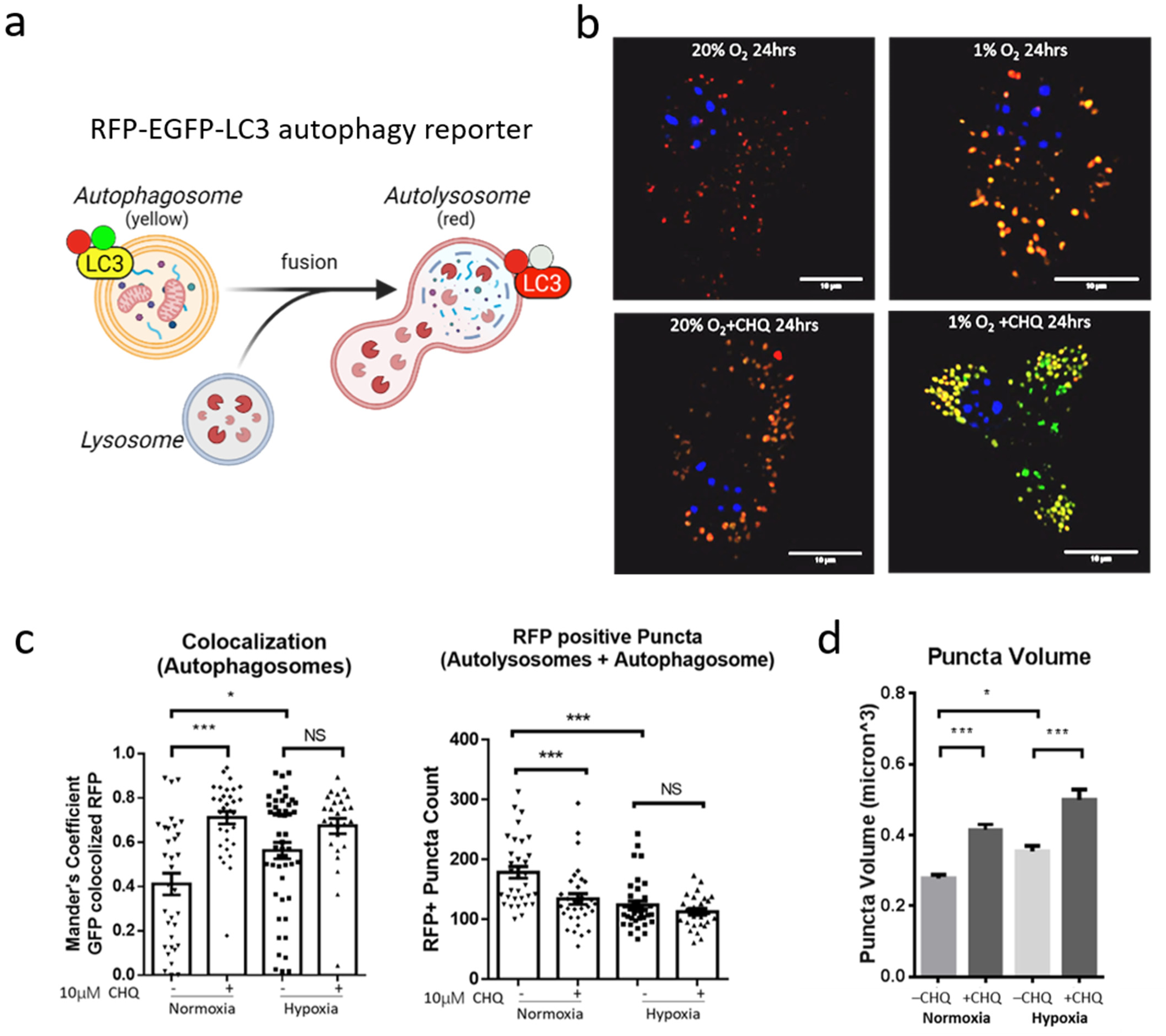
Prolonged hypoxia interrupts autophagic flux in mouse islet cells. Autophagic flux was analyzed by 3D confocal analysis of islet cells from CAG-RFP-EGFP-LC3 reporter mice, cultured for 24 hours under 20% O_2_ or 1% O_2_ with and without addition of 10 µM chloroquine (CHQ). a) Schematic illustrating the principles of the RFP-EGFP-LC3 fluorescent autophagy biosensor. RFP-EGFP-double-tagged LC3 accumulates on autophagosomes, which can be visualized as green and red (i.e. yellow) puncta. After fusion with a lysosome, EGFP fluorescence is quenched by the low pH in the lysosomal lumen, and the resulting autolysosome can be visualized as a red-only vesicle (created with BioRender.com). b) Representative 2D cross-sectional confocal images of RFP-EGFP-LC3-expressing islet cells showing autolysosomes in red (RFP single-positive) and autophagosomes and/or non-acidic autolysosomes in yellow (RFP/EGFP double-positive) under the various culture conditions. c) Colocalization of EGFP and RFP fluorescence was quantified on a per-cell basis using the Manders’ Overlap Coefficient, which ranges from 0 (no overlap) to 1 (complete overlap). Increased colocalization coefficients indicate autophagosome accumulation with preserved EGFP fluorescence in cells exposed to hypoxia and/or CHQ treatment. The combined number of autophagosomes and autolysosomes were quantified from the number of RFP-positive (RFP^+^) puncta in 3D reconstructions of individual cells. d) Quantification of individual RFP^+^ puncta shows that hypoxia- and CHQ-induced impairment of autophagic flux is associated with increased autophagosome/autolysosome volume. Data represent mean ± SEM of 3 independent experiments. \**p* < 0.05, \*\**p* < 0.01, \*\*\**p* < 0.001 by 2-way ANOVA followed by Tukey’s multiple comparison test.

### Hypoxia impairs the expression of β-cell Tfeb and its target genes

In agreement with hypoxia-induced dysregulation of lysosomes, we detected a reduction in islet expression of *Lamp1*, *Lamp2* and *Atp6v1h* (**Figure 5a**), which are involved in lysosomal membrane formation and luminal acidification [27]. The lysosomal mRNAs and autophagy-related transcripts that were reduced under hypoxic culture are all targets of TFEB [28–30]. We therefore examined *Tfeb* expression and found that prolonged hypoxia reduced *Tfeb* mRNA in mouse islets (**Figure 5b**) and decreased TFEB protein levels in both mouse and human β-cells (**Figures 5d,e**). To assess the role of Tfeb gene expression under hypoxia, we used an adenoviral vector (Ad.*Ins2*>mTFEB:T2A:TagBFP2) to increase Tfeb levels specifically in islet β-cells. Compared to islets transduced with control virus (Ad.*Ins2*>TagBFP2), β-cell overexpression of *Tfeb* successfully restored, and in some instances further amplified, the expression of lysosome-and autophagy-related Tfeb targets in islets cultured under 1% O_2_ (**Figure 5b**). Taken together, our data suggest that hypoxia may impair β-cell lysosomal function by disrupting Tfeb-dependent transcription of targets in the CLEAR gene network [31].

**Figure 5.**
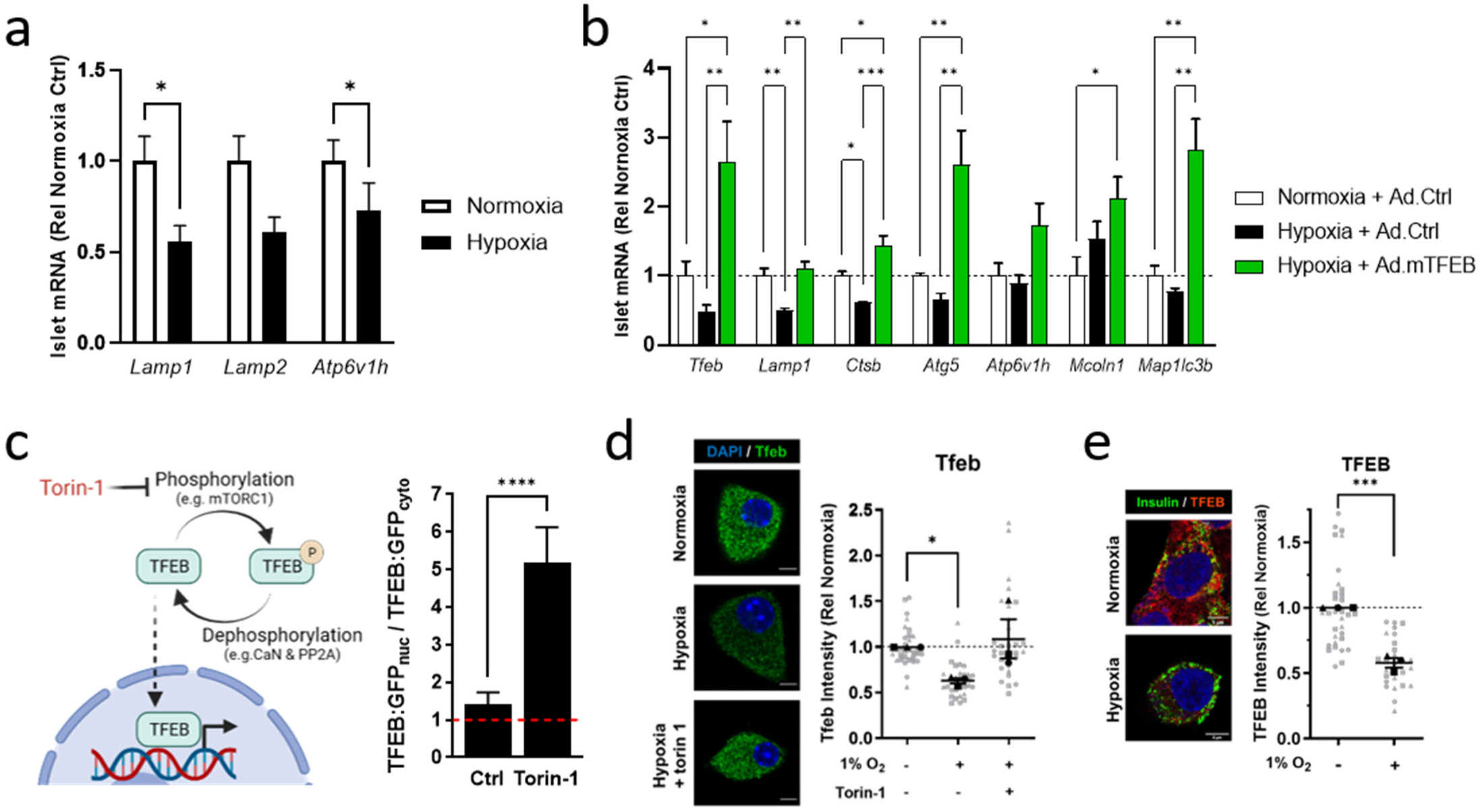
Hypoxic stress causes loss of β-cell Tfeb and Tfeb-regulated gene expression. a) qPCR analysis of changes to expression of lysosome-related genes in C57BL/6J islets cultured 16 hours under 20% O_2_ (normoxia) or 1% O_2_ (hypoxia) (n=6 independent experiments). b) qPCR analysis of *Tfeb* and select target genes related to lysosomes and autophagy in C57BL/6J islets cultured 16 hours under normoxia, or under hypoxia with and without β-cell-targeted adenoviral *Tfeb* overexpression. Data represent mean ± SEM of 4 independent experiments. c) *Left:* Schematic illustrating the role of phosphorylation and dephosphorylation on TFEB nuclear translocation and activation. Torin-1 inhibits mTORC1 and activates TFEB. Right: Quantified ratio of nuclear-to-cytosolic fluorescence in MIN6 cells transfected with plasmid expressing GFP-tagged TFEB. Data represent mean ± SEM of 42 Control and 29 torin-1 treated cells. d) *Left*: Representative Tfeb immunostaining. *Right*: Quantified Tfeb immunofluorescence intensity in male C57BL/6J β-cells cultured under 20% O_2_ and under 1% O_2_ with and without 500 nM torin-1 treatment for 24 hours. Grey markers represent the mean Tfeb fluorescence values of individual β-cells (identified by insulin staining, not shown). Black markers and lines show the mean ± SEM of 3 independent experiments. e) *Left*: Representative TFEB immunostaining. *Right*: Quantified TFEB immunofluorescence intensity in human β-cells cultured under 20% O_2_ and 1% O_2_ for 24 hours. Grey markers represent mean TFEB fluorescence values of individual β-cells (identified by insulin). Black markers and lines show the mean ± SEM of cells preparations from 3 non-diabetic donors. \**p* < 0.05, \*\**p* < 0.01, \*\*\**p* < 0.001 \*\*\*\**p* < 0.0001. Data in panels (a), (c) and (e) were analyzed by Student’s t-test. Data in panels (b) and (d) were analyzed by one-way ANOVA followed by Tukey’s multiple comparison test. In panels (a) and (b) each gene was analyzed separately.

### Inhibition of mTORC1 restores Tfeb, improves lysosomal function and reduces β-cell death under hypoxia

TFEB phosphorylation by mTOR inhibits its transcriptional activity by promoting its nuclear export and tethering to 14-3-3 proteins in the cytosol [32, 33]. Accordingly, TFEB nuclear translocation and transcriptional activity increase following mTORC1 inhibition by torin-1 (**Figure 5c**) [34]. Strikingly, we found that torin-1 prevented hypoxia-induced loss of Tfeb immunoreactivity in mouse β-cells (**Figure 5d**), possibly due to transcriptional autoregulation by residual Tfeb [35]. Torin-1-induced protection of Tfeb was paralleled by upregulation of several Tfeb target genes in hypoxic islets (**Figure 6a**). However, some of the transcripts we examined (*Lamp1*, *Lamp2* and *Atg5*) were resistant to this effect (**Figure 6a**) even if they were induced by *Tfeb* overexpression. The reason for this difference is unclear. Despite the reduction in islet *Lamp1* mRNA expression, vesicular Lamp1 immunoreactivity was maintained, and even tended to increase, indicating that β-cell lysosomes are not lost under hypoxia (**Figure 6b**). Lamp1 is an abundant lysosomal membrane protein, but it is not directly involved in the degradative function of the organelle. Islet expression of the lysosomal cathepsins, *Ctsb and Ctsd*, was reduced under hypoxia and could be restored by β-cell *Tfeb* overexpression and torin-1 treatment (**Figures 5b & 6a**). Moreover, hypoxia significantly impaired β-cell CTSB enzymatic activity, and this lysosomal defect was also partially ameliorated by torin-1 (**Figure 6c**). In the timeframe examined here, hypoxia-induced loss of β-cell Tfeb therefore negatively impacted lysosomal degradative function, but not the size of the lysosomal population.

**Figure 6.**
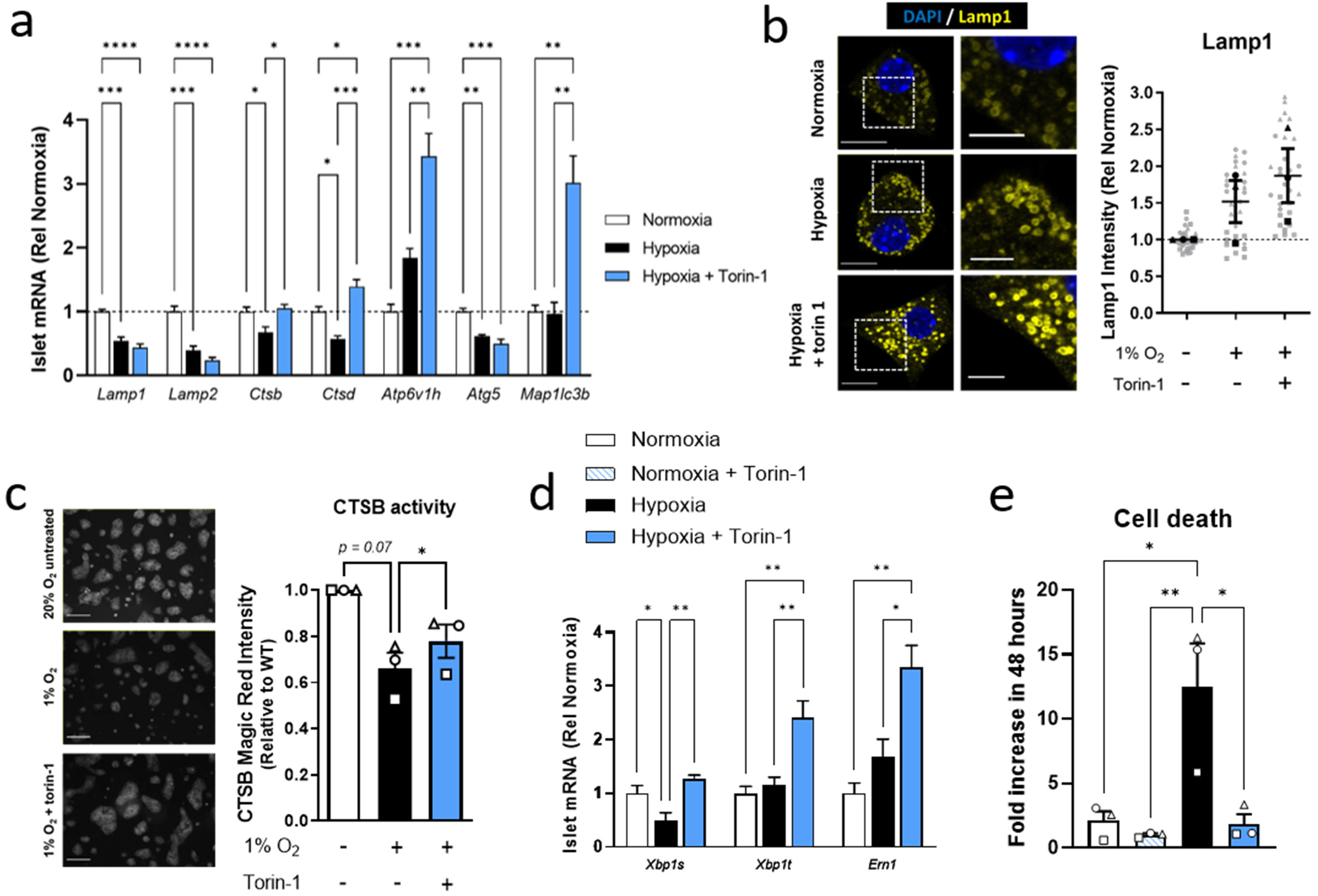
mTORC1 inhibition improves lysosomal function and β-cell survival under hypoxia. a) qPCR analysis of changes to expression of select TFEB target genes in C57BL/6J islets cultured 24 hours under 20% O_2_ (normoxia), or under 1% O_2_ (hypoxia) with and without 500 nM torin-1 (n=4 independent experiments). b) *Left:* Representative Lamp-1 immunostaining. Images in the right column show zooms of the areas indicated by white squares in the left column. *Right:* Quantified Lamp-1 immunofluorescence intensities. Grey markers represent the mean Lamp-1 intensity values of individual β-cells. Black markers and lines show the mean ± SEM of three independent experiments. Scale-bars = 10 μm in the original images and 5 μm in zoomed images. c) *Left:* Representative images showing Magic Red^®^ fluorescent staining of cathepsin B activity in C57BL/6J islet cells, and *Right*: quantified cathepsin B activity after 24 hours culture under 20% O_2,_ or under 1% O_2_ with and without 250 nM torin-1. Scale bars = 100 μm. d) qPCR quantification of UPR-related gene expression in C57BL/6J islets cultured under 20% O_2_ (normoxia) and 1% O_2_ (hypoxia) with and without 500 nM torin-1 for 16 hours. Data represent mean ± SEM of 4 independent experiments. e) Quantification of the fold increase in islet cell death during 48 hours culture under 20% O_2_ (normoxia), and 1% O_2_ (hypoxia) with and without 500 nM torin-1. Data represent mean ± SEM of 3 independent experiments. \**p* < 0.05, \*\**p* < 0.01, \*\*\**p* < 0.001 by one-way ANOVA followed by Tukey’s multiple comparison test. In panels (a) and (d) each gene was analyzed separately.

In addition to lysosomal autophagy, TFEB has been implicated in the regulation of cellular ER stress and integrated stress response pathways [36, 37]. We therefore re-examined the expression of UPR-related transcripts under hypoxia and found that torin-1 treatment significantly increased expression of *Ern1* (i.e. Ire1α) and normalized the levels of *Xbp1s* in hypoxic islets (**Figure 6d**). Finally, we asked whether the physiological benefits of mTORC1 inhibition enhanced resilience to hypoxia-induced stress and found that torin-1 treatment markedly protected mouse islet cells from hypoxia-induced cell death (**Figure 6e**).

Collectively, our data show that prolonged hypoxia impairs lysosomal function and disrupts protective autophagy in β-cells. This is linked to TFEB loss and reduced expression of its target genes, which can be restored through *Tfeb* overexpression or pharmacological activation. These findings highlight TFEB dysregulation as a contributor to the failure of lysosomal clearance mechanisms in hypoxic β-cells.

## DISCUSSION

In this study, we show that autophagy is an important adaptive process that protects pancreatic β-cells during hypoxia and following transplantation. Previous studies have established autophagy as a β-cell homeostatic pathway during metabolic and ER stress [20, 21], and recent work suggests that defective β-cell autophagy may also enhance islet immunogenicity [24], but its role during sustained oxygen deprivation and early graft stress has remained less clear. Our findings demonstrate that autophagy supports β-cell survival under hypoxia and further reveal that prolonged hypoxia compromises the lysosomal clearance capacity required for effective autophagic flux. This impairment was associated with reduced TFEB expression, decreased expression of lysosomal target genes, reduced cathepsin B activity, and accumulation of autophagosomes. Finally, we showed that pharmacologic inhibition of mTORC1 with torin-1 restored TFEB protein levels, improved lysosomal activity, and protected β-cells from hypoxia-induced death. Together, these findings support a model in which prolonged hypoxia impairs the autophagy–lysosome network and thereby compromises an important pathway for β-cell stress adaptation.

Syngeneic islet transplantation experiments extend the protective role of autophagy beyond isolated hypoxic culture. Short-term β-cell *Atg5* deletion did not detectably impair donor glucose tolerance or glucose-stimulated islet Ca^2+^ responses and insulin secretion before transplantation, but it accelerated marginal-mass graft failure. This distinction suggests that autophagy is not critical for basal β-cell function but becomes particularly important when β-cells encounter the combined stresses of engraftment. These data show that β-cell autophagy contributes to resilience during the early post-transplant stress period when grafts are exposed to hypoxia together with other metabolic and engraftment-associated stresses.

Our in vitro hypoxia experiments isolated one major component of the graft environment while allowing more direct analysis of autophagy-lysosome regulation. We demonstrated that hypoxia causes a marked impairment of autophagic flux and lysosomal function in β-cells. Accumulation of autophagosomes, increased autophagosome/autolysosome volume, and reduced cathepsin B activity collectively support the conclusion that prolonged oxygen limitation disrupts the late degradative phase of autophagy, rather than simply inducing formation of more autophagosomes. Because cathepsin B maturation and activity depend on the lysosomal environment, its reduced activity is consistent with impaired lysosomal function, potentially involving altered acidification, reduced cathepsin B maturation, or reduced enzyme abundance. Consistent with this functional impairment, hypoxia reduced the expression of autophagy- and lysosome-related genes, including *Atg5*, *Map1lc3b*, *Becn1*, *Lamp1*, *Ctsb*, and *Atp6v1h.* These findings identify lysosomal dysfunction as a point of failure in the β-cell response to sustained oxygen deprivation. The parallel reduction in several adaptive UPR transcripts further suggests that hypoxia weakens multiple proteostatic defense pathways, although the functional relationship between UPR impairment and lysosomal failure remains to be defined.

We further found that hypoxia reduced TFEB expression at both mRNA and protein levels. This was accompanied by decreased expression of TFEB-dependent lysosomal genes and reduced lysosomal enzyme activity, pointing to TFEB loss as a potential link between oxygen limitation and impaired lysosomal function. In contrast to nutrient-replete conditions, where mTORC1-dependent phosphorylation retains TFEB in the cytoplasm without necessarily reducing TFEB expression [32, 33], hypoxia reduced overall TFEB abundance. Because TFEB coordinates lysosomal and autophagic gene networks [28–31], loss of TFEB likely contributes to the observed impairment of autophagic flux and lysosomal activity. The observation that TFEB protein also declined in human β-cells further supports the relevance of these findings. However, additional work will be needed to determine whether hypoxia similarly impairs lysosomal autophagy in human β-cells and if restoring TFEB can improve their resilience to hypoxic stress.

The mechanism by which hypoxia reduces TFEB levels remains unclear. Both Tfeb mRNA and protein declined under hypoxia, indicating that reduced gene expression likely contributes, but effects on translation remain possible. For example, Programmed Cell Death 4 (PDCD4) can suppress TFEB translation in other cell types [38], and PDCD4 is induced by hypoxia in MIN6 β-cells [39], suggesting that multiple levels of control may play a role in hypoxic β-cells. TFEB activity is also modulated post-translationally through phosphorylation-dependent control of its subcellular localization, most notably by mTORC1 [32, 33], and may also be influenced by AMPK-linked energy-sensing pathways [40]. Although we did not assess TFEB phosphorylation or localization in this study, these pathways are likely candidates for mediating crosstalk between oxygen availability and lysosomal adaptation. Future studies using targeted manipulation and characterization of these regulators may help define how hypoxia affects TFEB activation and turnover in β-cells.

Under nutrient-replete conditions, mTORC1 phosphorylates TFEB, retaining it in the cytoplasm and thereby suppressing its transcriptional program without directly reducing TFEB transcription [32, 33]. In hypoxic β-cells, pharmacologic inhibition of mTORC1 with torin-1 restored the expression of TFEB and several downstream genes, improved lysosomal activity and cell viability. This is consistent with relief of mTORC1-dependent inhibition of TFEB, which may allow reinforcement of transcriptional feedback, whereby nuclear TFEB binds its own promoter to maintain expression of itself and its targets [35]. Our torin-1 results demonstrate that the TFEB-lysosomal axis remains modifiable during hypoxic stress. However, because torin-1 has TFEB-independent effects and mTORC1 activity was not directly measured, we cannot conclude that mTORC1 is the primary upstream regulator of TFEB under hypoxia. Nonetheless, these findings highlight the potential for oxygen and nutrient signaling to converge on TFEB regulation and provide a framework for future mechanistic testing.

The impairment of autophagy and lysosomal function under hypoxia has implications for β-cell biology in both transplantation and diabetes. In islet transplantation, early graft hypoxia is a major cause of β-cell loss before revascularization. Strategies to reduce early graft loss include efforts to improve oxygen delivery, vascularization, immune protection, or transplant-site design [5]. Our findings that autophagy is protective, and that its failure coincides with reduced TFEB expression, complement these approaches and suggest that pharmacologic or genetic strategies that maintain TFEB activity or lysosomal competence may help mitigate early graft loss. However, because TFEB and mTORC1 can also influence insulin biosynthesis and secretion [41, 42], therapeutic manipulation should carefully balance enhanced stress resilience and potential effects on β-cell function. In diabetes, where islet hypoxia may arise from reduced vascular support and high metabolic demand [43, 44], similar pathways may contribute to progressive β-cell decline. Defining how oxygen limitation alters TFEB and lysosomal regulation could thus inform strategies to preserve β-cell mass in both transplantation and metabolic disease.

In summary, we demonstrated that autophagy is protective for β-cells during hypoxia and early transplantation-induced stress, but that sustained hypoxia impairs this defense by disrupting TFEB-associated gene expression and lysosomal function. The accompanying loss of autophagic flux may contribute to reduced β-cell survival and graft vulnerability. Pharmacologic restoration of TFEB with torin-1 ameliorated hypoxia-induced defects in vitro, consistent with the idea that mTORC1-dependent inhibition of TFEB limits lysosomal capacity during hypoxic stress. Although the upstream signals linking hypoxia to TFEB and autophagy suppression remain to be defined, our findings establish a framework connecting oxygen availability, lysosomal function, and β-cell survival in settings relevant to both islet transplantation and diabetes.

## ACKNOWLEDGEMENTS

This work was funded by a Canucks for Kids Diabetes Fund Catalyst Grant and a Breakthrough T1D (formerly JDRF) award to D.S.L. (CDA 2-2013-50). D.S.L. received salary support from BC Children’s Hospital Research Institute (BCCHR). Y.Z. was supported by a Doctoral Studentship Award from BCCHR. We thank Dr P Lavoie (BCCHR) for use of their hypoxic chamber and Dr F Lynn (BCCHR) for sharing adenoviral constructs for Cre expression. Finally, we acknowledge expert technical assistance from Benny Tang (BCCHR) and Dr. Jingsong Wang (BCCHR Imaging Core; RRID:SCR_026573) and technical support from the In Vitro Phenotyping Core of the Breakthrough T1D Canucks for Kids Centre of Excellence at UBC.

The authors acknowledge that UBC and BC Children’s Hospital are situated on the traditional, ancestral, and unceded territories of the S wxwú7mesh (Squamish), Sәlı’lwәta /Selilwitulh (Tsleil-Waututh), and x mәqk әyәm (Musqueam) Nations.

## AUTHOR CONTRIBUTIONS

Y.Z. and D.S.L. designed the study and wrote the paper. Y.Z., D.J.P., R.T., M.K., D.L.D., G.S. and D.S.L. performed experiments and analyzed data. Y.Z., D.J.P., C.B.V. and D.S.L interpreted data and edited the paper.

## REFERENCES

[1] Gerber PA, Rutter GA (2017) The Role of Oxidative Stress and Hypoxia in Pancreatic Beta-Cell Dysfunction in Diabetes Mellitus. Antioxid Redox Signal 26(10): 501–518. 10.1089/ars.2016.6755

[2] Carlsson PO, Liss P, Andersson A, Jansson L (1998) Measurements of oxygen tension in native and transplanted rat pancreatic islets. Diabetes 47(7): 1027–1032. 10.2337/diabetes.47.7.1027

[3] Lau J, Henriksnas J, Svensson J, Carlsson PO (2009) Oxygenation of islets and its role in transplantation. Curr Opin Organ Transplant 14(6): 688–693. 10.1097/MOT.0b013e32833239ff

[4] Olsson R, Olerud J, Pettersson U, Carlsson PO (2011) Increased numbers of low-oxygenated pancreatic islets after intraportal islet transplantation. Diabetes 60(9): 2350–2353. 10.2337/db09-0490

[5] Gamble A, Pepper AR, Bruni A, Shapiro AMJ (2018) The journey of islet cell transplantation and future development. Islets 10(2): 80–94. 10.1080/19382014.2018.1428511

[6] Faleo G, Russ HA, Wisel S, et al. (2017) Mitigating Ischemic Injury of Stem Cell-Derived Insulin-Producing Cells after Transplant. Stem Cell Reports 9(3): 807–819. 10.1016/j.stemcr.2017.07.012

[7] Marasco MR, Linnemann AK (2018) beta-Cell Autophagy in Diabetes Pathogenesis. Endocrinology 159(5): 2127–2141. 10.1210/en.2017-03273

[8] Mazure NM, Pouyssegur J (2010) Hypoxia-induced autophagy: cell death or cell survival? Curr Opin Cell Biol 22(2): 177–180. 10.1016/j.ceb.2009.11.015

[9] Song S, Tan J, Miao Y, Sun Z, Zhang Q (2018) Intermittent-Hypoxia-Induced Autophagy Activation Through the ER-Stress-Related PERK/eIF2alpha/ATF4 Pathway is a Protective Response to Pancreatic beta-Cell Apoptosis. Cell Physiol Biochem 51(6): 2955–2971. 10.1159/000496047

[10] Hara T, Nakamura K, Matsui M, et al. (2006) Suppression of basal autophagy in neural cells causes neurodegenerative disease in mice. Nature 441(7095): 885–889. 10.1038/nature04724

[11] Aharoni-Simon M, Shumiatcher R, Yeung A, et al. (2016) Bcl-2 Regulates Reactive Oxygen Species Signaling and a Redox-Sensitive Mitochondrial Proton Leak in Mouse Pancreatic beta-Cells. Endocrinology 157(6): 2270–2281. 10.1210/en.2015-1964

[12] Luciani DS, White SA, Widenmaier SB, et al. (2013) Bcl-2 and Bcl-xL Suppress Glucose Signaling in Pancreatic beta-Cells. Diabetes 62(1): 170–182. 10.2337/db11-1464

[13] Courtade JA, Wang EY, Yen P, et al. (2017) Loss of prohormone convertase 2 promotes beta cell dysfunction in a rodent transplant model expressing human pro-islet amyloid polypeptide. Diabetologia 60(3): 453–463. 10.1007/s00125-016-4174-2

[14] Montane J, Bischoff L, Soukhatcheva G, et al. (2011) Prevention of murine autoimmune diabetes by CCL22-mediated Treg recruitment to the pancreatic islets. J Clin Invest 121(8): 3024–3028. 10.1172/JCI43048

[15] Sabatini PV, Speckmann T, Nian C, et al. (2018) Neuronal PAS Domain Protein 4 Suppression of Oxygen Sensing Optimizes Metabolism during Excitation of Neuroendocrine Cells. Cell Reports 22(1): 163–174. 10.1016/j.celrep.2017.12.033

[16] Wang X, Hao J, Metzger DL, et al. (2009) Local expression of B7-H4 by recombinant adenovirus transduction in mouse islets prolongs allograft survival. Transplantation 87(4): 482–490. 10.1097/TP.0b013e318195e5fa00007890-200902270-00004 [pii]

[17] Hammar E, Parnaud G, Bosco D, et al. (2004) Extracellular matrix protects pancreatic beta-cells against apoptosis: role of short- and long-term signaling pathways. Diabetes 53(8): 2034–2041. 10.2337/diabetes.53.8.2034

[18] Bensellam M, Maxwell EL, Chan JY, et al. (2016) Hypoxia reduces ER-to-Golgi protein trafficking and increases cell death by inhibiting the adaptive unfolded protein response in mouse beta cells. Diabetologia 59(7): 1492–1502. 10.1007/s00125-016-3947-y

[19] Choi SE, Lee SM, Lee YJ, et al. (2009) Protective role of autophagy in palmitate-induced INS-1 beta-cell death. Endocrinology 150(1): 126–134. 10.1210/en.2008-0483

[20] Ebato C, Uchida T, Arakawa M, et al. (2008) Autophagy is important in islet homeostasis and compensatory increase of beta cell mass in response to high-fat diet. Cell Metab 8(4): 325–332. S1550-4131(08)00250-7 [pii] 10.1016/j.cmet.2008.08.009

[21] Bachar-Wikstrom E, Wikstrom JD, Kaiser N, Cerasi E, Leibowitz G (2013) Improvement of ER stress-induced diabetes by stimulating autophagy. Autophagy 9(4): 626–628. 10.4161/auto.23642

[22] Kuma A, Hatano M, Matsui M, et al. (2004) The role of autophagy during the early neonatal starvation period. Nature 432(7020): 1032–1036. 10.1038/nature03029

[23] Jung HS, Chung KW, Won Kim J, et al. (2008) Loss of autophagy diminishes pancreatic beta cell mass and function with resultant hyperglycemia. Cell Metab 8(4): 318–324. S1550-4131(08)00254-4 [pii] 10.1016/j.cmet.2008.08.013

[24] Austin MC, Muralidharan C, Roy S, Crowder JJ, Piganelli JD, Linnemann AK (2025) Dysfunctional beta-cell autophagy induces beta-cell stress and enhances islet immunogenicity. Front Immunol 16: 1504583. 10.3389/fimmu.2025.1504583

[25] Pei X, Wang H, Xu P, Liang K, Yuan L (2022) The core autophagy protein ATG5 controls the polarity of the Golgi apparatus and insulin secretion of pancreatic beta cells. Biochem Biophys Res Commun 629: 26–33. 10.1016/j.bbrc.2022.08.084

[26] Li L, Wang ZV, Hill JA, Lin F (2014) New autophagy reporter mice reveal dynamics of proximal tubular autophagy. J Am Soc Nephrol 25(2): 305–315. 10.1681/ASN.2013040374

[27] Huynh KK, Eskelinen EL, Scott CC, Malevanets A, Saftig P, Grinstein S (2007) LAMP proteins are required for fusion of lysosomes with phagosomes. EMBO J 26(2): 313–324. 10.1038/sj.emboj.7601511

[28] Palmieri M, Impey S, Kang H, et al. (2011) Characterization of the CLEAR network reveals an integrated control of cellular clearance pathways. Human molecular genetics 20(19): 3852–3866. 10.1093/hmg/ddr306

[29] Settembre C, Di Malta C, Polito VA, et al. (2011) TFEB Links Autophagy to Lysosomal Biogenesis. Science. science.1204592 [pii] 10.1126/science.1204592

[30] Sardiello M, Palmieri M, di Ronza A, et al. (2009) A gene network regulating lysosomal biogenesis and function. Science 325(5939): 473–477. 10.1126/science.1174447

[31] Settembre C, Medina DL (2015) TFEB and the CLEAR network. Methods Cell Biol 126: 45–62. 10.1016/bs.mcb.2014.11.011

[32] Martina JA, Chen Y, Gucek M, Puertollano R (2012) MTORC1 functions as a transcriptional regulator of autophagy by preventing nuclear transport of TFEB. Autophagy 8(6): 903–914. 10.4161/auto.19653

[33] Napolitano G, Esposito A, Choi H, et al. (2018) mTOR-dependent phosphorylation controls TFEB nuclear export. Nature communications 9(1): 3312. 10.1038/s41467-018-05862-6

[34] Settembre C, Zoncu R, Medina DL, et al. (2012) A lysosome-to-nucleus signalling mechanism senses and regulates the lysosome via mTOR and TFEB. EMBO J 31(5): 1095–1108. 10.1038/emboj.2012.32

35. Settembre C, De Cegli R, Mansueto G, et al. (2013) TFEB controls cellular lipid metabolism through a starvation-induced autoregulatory loop. Nat Cell Biol 15(6): 647–658. 10.1038/ncb2718

[36] Martina JA, Diab HI, Brady OA, Puertollano R (2016) TFEB and TFE3 are novel components of the integrated stress response. EMBO J 35(5): 479–495. 10.15252/embj.201593428

[37] Zhang Z, Qian Q, Li M, et al. (2020) The unfolded protein response regulates hepatic autophagy by sXBP1-mediated activation of TFEB. Autophagy: 1–15. 10.1080/15548627.2020.1788889

[38] Cao B, Chen X, Li Y, et al. (2024) PDCD4 triggers alpha-synuclein accumulation and motor deficits via co-suppressing TFE3 and TFEB translation in a model of Parkinson’s disease. NPJ Parkinsons Dis 10(1): 146. 10.1038/s41531-024-00760-9

[39] Kumar S, Marriott CE, Alhasawi NF, Bone AJ, Macfarlane WM (2017) The role of tumour suppressor PDCD4 in beta cell death in hypoxia. PLoS One 12(7): e0181235. 10.1371/journal.pone.0181235

[40] Malik N, Ferreira BI, Hollstein PE, et al. (2023) Induction of lysosomal and mitochondrial biogenesis by AMPK phosphorylation of FNIP1. Science 380(6642): eabj5559. 10.1126/science.abj5559

[41] Pasquier A, Pastore N, D’Orsi L, et al. (2023) TFEB and TFE3 control glucose homeostasis by regulating insulin gene expression. EMBO J: e113928. 10.15252/embj.2023113928

[42] Israeli T, Riahi Y, Garzon P, et al. (2022) Nutrient Sensor mTORC1 Regulates Insulin Secretion by Modulating beta-Cell Autophagy. Diabetes 71(3): 453–469. 10.2337/db21-0281

[43] Sato Y, Endo H, Okuyama H, et al. (2011) Cellular hypoxia of pancreatic {beta}-cells due to high levels of oxygen consumption for insulin secretion in vitro. J Biol Chem. M110.194738 [pii] 10.1074/jbc.M110.194738

[44] Okajima Y, Matsuzaka T, Miyazaki S, et al. (2022) Morphological and functional adaptation of pancreatic islet blood vessels to insulin resistance is impaired in diabetic db/db mice. Biochim Biophys Acta Mol Basis Dis 1868(4): 166339. 10.1016/j.bbadis.2022.166339

