## Supplemental Figures for "Autophagy protects pancreatic β-cells during hypoxia and islet transplantation but is compromised by TFEB-lysosomal dysfunction"

**Zou et al. - Supplemental Figure 1**

a

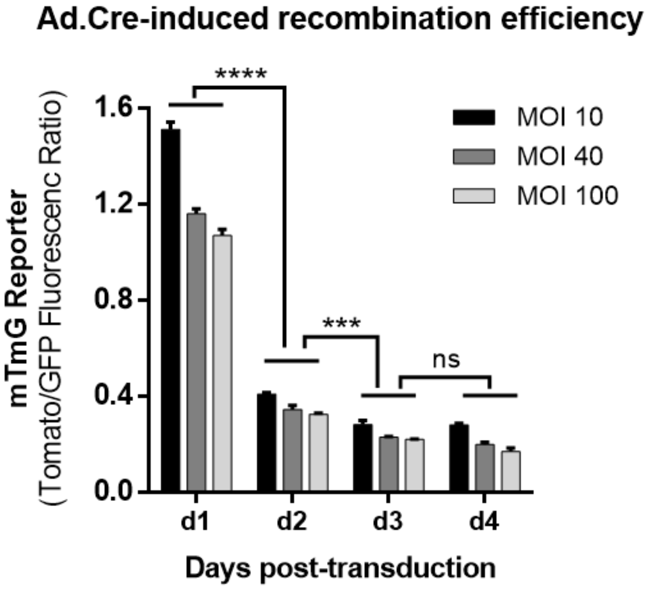


b

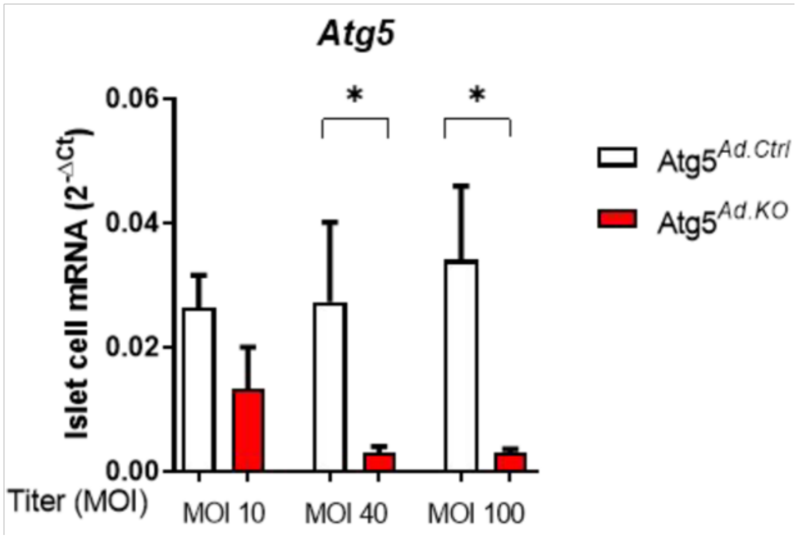


**a)** The ratio of tdTomato/EGFP reporter fluorescence intensities was quantified and used as an indicator of recombination over time at different MOIs of virus. ****$p < 0.0001$ by two-way ANOVA followed by Tukey's multiple comparison test.

**b**) *Atg5* mRNA levels in *Atg5*$^{flox/flox}$ mouse islet 4 days after transduction with 40 MOI Ad.Cre virus (*Atg5*$^{Ad.KO}$ cells) or Ad.Ctrl virus (*Atg5*$^{Ad.Ctrl}$ cells) n = 3; *$p < 0.05$ by paired Student's t-test.

### Zou et al. - Supplemental Figure 2

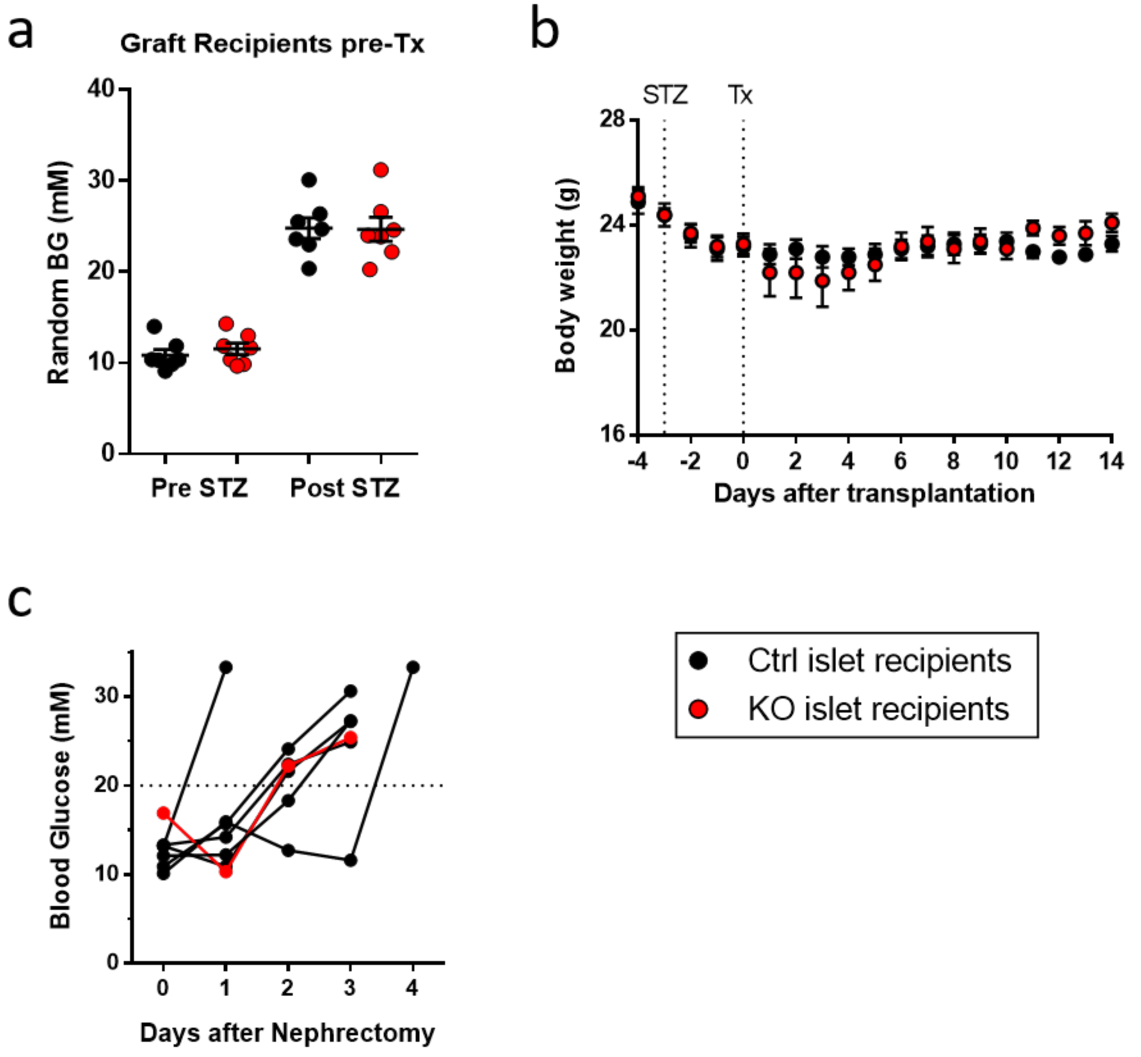


**a)** Blood glucose of recipient mice before and after STZ-induced diabetes but before islet transplantation (Tx).

**b**) Body weight of recipient mice before and after Tx.

**c)** Blood glucose levels of normoglycemic Tx recipients after nephrectomy confirming that glucose homeostasis at endpoint depended on islet graft.
